# A Critical Review on the Use of Support Values in Tree Viewers and Bioinformatics Toolkits

**DOI:** 10.1101/035360

**Authors:** Lucas Czech, Jaime Huerta-Cepas, Alexandros Stamatakis

**Affiliations:** Heidelberg Institute for Theoretical Studies, Schloss-Wolfsbrunnenweg 35, 69118 Heidelberg, Germany.; Structural and Computational Biology Unit, European Molecular Biology Laboratory, Meyerhofstrasse 1, 69117 Heidelberg, Germany.; Karlsruhe Institute of Technology, Institute for Theoretical Informatics, Am Fasanengarten 5, 76131 Karlsruhe, Germany.

**Keywords:** Phylogenetic trees, Tree visualization, Tree viewers, Bioinformatics toolkits, Newick format, Branch support values, Branch labels, Software, Bugs

## Abstract

Phylogenetic trees are routinely visualized to present and interpret the evolutionary relationships of species. Virtually all empirical evolutionary data studies contain a visualization of the inferred tree with branch support values. Ambiguous semantics in tree file formats can lead to erroneous tree visualizations and therefore to incorrect interpretations of phylogenetic analyses.

Here, we discuss problems that can and do arise when displaying branch values on trees after re-rooting. Branch values are typically stored as node labels in the widely-used Newick tree format. However, such values are attributes of branches. Storing them as node labels can therefore yield errors when re-rooting trees. This depends on the mostly implicit semantics that tools deploy to interpret node labels.

We reviewed 10 tree viewers and 10 bioinformatics toolkits that can display and re-root trees. We found that 14 out of 20 of these tools do not permit users to select the semantics of node labels. Thus, unaware users might obtain incorrect results when rooting trees inferred by common phylogenetic inference programs. We illustrate such incorrect mappings for several test cases and real examples taken from the literature. This review has already led to improvements and workarounds in 8 of the tested tools. We suggest tools should provide an option that explicitly forces users to define the semantics of node labels.

## I. Introduction

### A. Problem Description

The Newick format is widely used to store and visualize phylogenies. Archie et al. introduced it in 1986 [1]. Since then, it has become the de-facto standard for storing, exchanging, and displaying phylogenies. It uses parentheses and commas to specify the nesting structure of the tree and also allows for storing node labels as well as branch lengths.

In many cases, additional vital information needs to be associated with the branches of a tree. Published phylogenies usually display branch values, such as boot-strap [2] support, Bayesian posterior probability [3], or aLRT test [4] values. These values are associated with branches (splits/bipartitions) of the tree and not nodes of the tree. In the original specification of the Newick format, the authors had not foreseen an option for specifying branch labels or other meta-data associated to branches.

Thus, as a workaround, branch values are often stored as inner node labels in the output of phylogenetic inference tools. Node labels of tip nodes usually contain the species name of the extant organisms. Inner nodes, however, represent hypothetical common ancestors and are therefore generally not named. Thus, these inner node labels can be (mis-)used to store branch information.

The original Newick format is well-defined, for example via the formal grammar provided by [5]. There is however no official standard for it, including respective semantics of Newick comments, for instance. Hence, there is also no officially correct way of using it—attributes of branches and nodes can essentially be interpreted ad libitum. Thus, users need to be aware of the semantics of such attributes. Their interpretation depends on the convention used when storing those values in Newick format.

The convention, or rather workaround, for storing branch values as node labels exhibits potential pitfalls. This is because, in an unrooted binary tree, it is not clear to which of the three outgoing branches of an inner node such a node label refers to.

However, for rooted trees, there is an unambiguous mapping of node labels to branches: The node label (branch value) at an inner node can always be associated with (or mapped to) the outgoing branch that points toward the root. Note that, unrooted trees often have a dedicated inner node that serves as a hook for both computing and visualizing the tree. This so called top-level trifurcation is not a root in the strict sense, but required for storing and parsing the tree, because we need to recursively start traversing the tree from somewhere. We can choose the inner node that serves as top-level trifurcation arbitrarily. That is, the same underlying unrooted tree can be displayed or written to file in many distinct ways. For *n* taxa, an unrooted binary tree has n − 2 inner nodes, hence we can choose n − 2 such top-level trifurcations. For each of these possible top-level trifurcations, we can then also freely chose by which order we descend into the sub-trees defined by the three outgoing branches to print out or visualize the tree. The chosen top-level trifurcation induces an artificial orientation for branches in the tree, and can thus be used to unambiguously associate node labels with branches. Figure 1(a) shows an unrooted tree with this structure.

**Figure 1:**
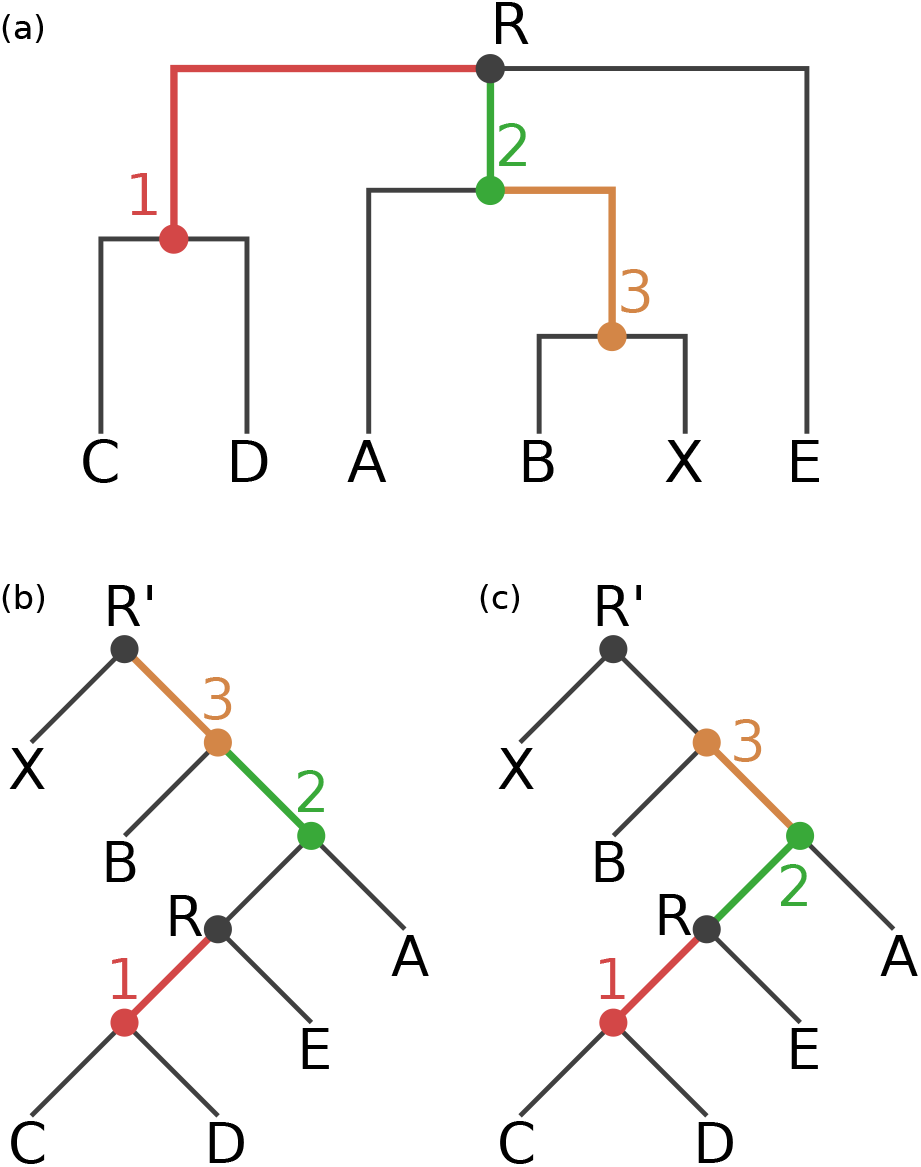
Our exemplary tree, before and after rooting on the branch leading to the tip node `X`. The rooted trees contain an additional root node R′. (a) Original rooting (via top-level trifurcation) and visual representation of our Newick test tree *T_N_*. Inner nodes and branches are colored according to the correct node label to branch mapping of *T_N_*. (b) Tree rooted on node R′. Node labels are mapped incorrectly to branches, resulting in a tree with an erroneous node label to branch value mapping. (c) Tree rooted on node `R`′. Node labels are correctly mapped to the branches of the tree.

Thus, both rooted and unrooted trees in Newick format explicitly (root) or implicitly (top-level trifurcation) encode a direction for branches. Therefore, the mapping between branch values and node labels in Newick files is well defined in principle: For restoring the correct association between node labels and branches, the direction towards the top-level node (root or top-level trifurcation) can be used. This however entails an implicit semantic interpretation. When reading a Newick-formatted tree, the user or program needs to know if inner node labels need to be interpreted as branch values. When this semantic distinction is not made, node labels need to be interpreted as being associated to the nodes, because this was the original intention of the Newick format. When node labels that should be interpreted as branch labels (e.g., support values) are erroneously interpreted as node labels, this can lead to incorrect visualizations as well as interpretations of phylogenies. These issues can potentially affect downstream analysis tools that parse phylogenies with node labels, for instance, tools for computing the weighted Robinson-Foulds distance [6] between phylogenies with branch support values.

Here, we show that 14 out of 20 common tree viewers and general purpose bioinformatics toolkits do not offer an explicit option for specifying the semantics of inner node labels. A simple way to examine the behavior of tools in this regard, is to (re-)root a given tree—an option that all tested viewers and toolkits offer. If node labels shall represent branch labels, the association of some node labels with corresponding branches has to be changed during the re-rooting process. This is because the direction towards the root (or top-level trifurcation) changes. We found that, 8 out of 20 tools exhibit incorrect behavior when re-rooting trees.

Note that, re-rooting a tree is not always a meaningful operation. For example, a tree inferred with a time-asymmetric model might contain posterior support values that belong to nodes rather than branches/splits of the tree. As another example, inner node labels that represent clade names (e.g., “Mammalia”) are attributes associated with one direction of a branch (only mammals in one part of the split induced by the branch, none in the other). In fact, this is a third class of values associated with the tree, which, again, behaves differently when re-rooting the tree. We are, however, not aware of any tree file format that allows to store this type of information. Thus, we focus on the distinction between node labels and branch values here, and use re-rooting to reveal the internal workings of the tested tools.

### B. Test Case

Our unrooted bifurcating Newick test tree with inner node labels

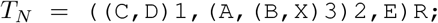

has six leaf nodes (`A`…`E`) and four inner nodes (labeled 1…3, and the top-level trifurcation `R`). For the sake of simplicity, we ignore branch length values. We use T_N_ throughout this review to test the behavior of tree viewers and toolkits when re-rooting the original topology. We also outline potential problems that may arise due to the mostly implicit semantics of inner node labels in Newick trees.

An alternative variant to output branch values is to store them as Newick comments in square brackets instead of node labels. The tree

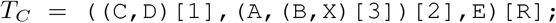

shows an example for this notation and contains the same information as tree *T_N_*. For the semantics and the association of those comments with branches, the same convention applies as for the node label notation. Some of the tested tools are able to correctly parse and display this format, but, in general, the same semantic issues and mapping problems arise.

For example, the output formats for phylogenies with branch support values of three widely-used phylogenetic inference tools are different. **PHYML** [7] reports support values as node labels, see [8]. **RAxML** [9] generates two tree files, one with comments and one with node labels. Finally, **MrBayes** [10], [11] uses its own Nexus-based format, which internally uses a variation of Newick comments to report support values (posterior probabilities). Those different idiosyncratic output formats illustrate the difficulties associated to working with trees that have branch support values.

In Figure 1 we show tree *T_N_*, where colors indicate the correct mapping of inner node labels to nodes and branches.

If we now (re-)root *T_N_* at the branch that leads to tip `X`, the mappings between all nodes and branches that lie on the path between the old and the new top-level node have to be altered. In our example, the nodes on the path from `R` to `X` are the inner nodes 2 and 3. In Figure 1, we display the incorrect (1(b)) and correct (1(c)) mapping of inner node labels to nodes and branches after re-rooting. Note that, this rooted binary tree now contains one more node, which is the newly created root node R'. In both Figures, the inner node labels are correctly assigned to their corresponding nodes. However, the association of those labels to the corresponding branches is only correct in Figure 1(c).

An incorrect mapping of node labels to branches as presented in Figure 1(b) will lead to incorrectly displayed branch values in empirical phylogenetic studies. In addition, since a typically large fraction of the results and discussion sections of such studies is dedicated to interpreting the support values of the phylogeny, the conclusions of these studies might also be incorrect.

In the following, we examine different popular tree viewers and several bioinformatics toolkits to determine if they maintain the correct branch value mapping when re-rooting our test tree TN at the branch leading to tip node X.

Finally, since Dendroscope [12], one of the most commonly used tree viewers tested, yielded incorrect mappings for all versions prior to v. 3.5.0 (released 2016-01-07), we also assessed if there exist published empirical phylogenetic studies using Dendroscope with incorrectly visualized support values.

## II. Review

### A. Experimental Setup

Given a Newick tree with inner node labels (e.g., tree *T_N_* with labels 1, 2 and 3), we distinguish between two possible interpretations for those labels: i) They are actual node labels (e.g., ancestral species names). We call this the “node interpretation”, and ii) they represent branch labels (e.g., support values). We call this the “branch interpretation”. The same applies to trees that use comments instead of node labels (e.g., tree *T_C_*). For a program to support both interpretations, a reasonable solution would be to offer an option for choosing between the two, that is, to include an explicit semantic interpretation dialog.

We tested the tree viewers as follows:

- Check whether the tool has an option to specify the semantics of inner values.
- Load trees *_N_* and *T_C_* from the corresponding Newick file.
- Check how the tool interprets the values.
- Re-root the tree at the branch leading to node `X`.
- Check whether the viewer works correctly based on its interpretation.

In Table I, we provide an overview of the tested tree viewers and bioinformatics toolkits. While the list does not cover all available tools, we focus on highly used resources offering re-rooting capabilities, as the impact of potential errors in these tools on published phylogenies is larger. We also tested some less known tools, in order to assess how widely spread the issue is.

**Table I:** Evaluated tree viewers and bioinformatics toolkits with accumulated number of citations (https://scholar.google.com, accessed on 2016-11-11).

| Tool | Version | Reference | Citations |
| --- | --- | --- | --- |
| Archaeopteryx | 0.9911 | [13] | 268 |
| ATV | 4.00 alpha 13 | [14] | 288 |
| Dendroscope | 3.4.0 and 3.5.3 | [12] | 1,348 |
| ETE (GUI) | 2.3.10 | [15] | 238 |
| EvolView | Accessed 2016-08-15 | [16] | 105 |
| FigTree | 1.4.2 | [17] | > 2,362* |
| iTOL | Accessed 2016-08-15 | [18] | 1,879 |
| PhyloWidget | Accessed 2016-08-15 | [19] | 113 |
| TreeView | 1.6.6 (Windows) | [20] | 10,570 |
| T-REX | Accessed 2016-08-15 | [21] | 285 |
| APE | 3.4 | [22] | 3,915 |
| BioPerl | 1.006925 | [23] | 1,410 |
| BioPython | 1.63b | [24] | 797 |
| Dendropy | 4.1.0 | [25] | 525 |
| ETE (API) | 3.0.0b35 | [15] | 238 |
| Geneious | 10.0.5 | [26] | 1,689 |
| Mega | 7.0.14 build 7160126 | [27] | 69,134 |
| Mesquite | 3.10 (build 765) | [28] | 5,616 |
| Newick Utilities | 1.6 | [29] | 31 |
| Pycogent/scikit-bio | 1.5.3 | [30] | 148 |
| Total |  |  | 100,721 |

\* FigTree does not have an official publication, so we estimated the number of citations by accumulating the counts for the most recent versions.

In the following, we discuss our observations for the aforementioned tree viewers and general purpose toolkits. In Table II we provide an overview of these results.

**Table II:** Evaluation of tree viewers and bioinformatics toolkits. The columns “Nodes” and “Branches” indicate which of the two interpretations of Newick node labels the tool supports. The last column shows whether the re-rooting behavior is correct according to the interpretation offered or implied by the tool.

| Tool | Nodes | Branches | Default behavior | Correct re-rooting |
| --- | --- | --- | --- | --- |
| Archaeopteryx | ✓ | ✓ | nodes | ✓ |
| ATV |  | ✓ | branches | ✓ |
| Dendroscope | ✓ | ✓* | dialog* | ✓* |
| ETE (GUI) | ✓ | ✓ | branches | ✓ |
| EvolView |  | ✓ | branches |  |
| FigTree | ✓ | ✓ | both |  |
| iTOL | ✓ | ✓ | input dependent | ✓ |
| PhyloWidget | ✓ |  | nodes |  |
| TreeView |  | ✓ | branches | ✓ |
| T-REX |  | ✓ | branches | (✓) |
| APE | ✓ | ✓* | nodes | ✓* |
| BioPerl | ✓ | ✓ | nodes | ✓ |
| BioPython | ✓ |  | nodes |  |
| Dendropy | ✓ | ✓* | nodes | (✓) |
| ETE (API) | ✓ | ✓ | branches | ✓ |
| Geneious | ✓ | (✓) | nodes |  |
| MEGA |  | ✓ | branches | ✓ |
| Mesquite | ✓ | ✓ | nodes | ✓ |
| Newick Utilities | ✓ | ✓* | nodes | ✓* |
| Pycogent/scikit-bio |  | ✓ | branches | ✓ |

\* Option added or improved after this review.

### B. Results

#### 1 Tree Viewers

**Archaeopteryx** is aware of the semantic issue, see [31]. It offers an option to define the semantics of annotated values. The default is to interpret nodes labels as node labels, thus the re-rooted tree is correctly displayed only for the node interpretation. When activating the option “Internal Node Names are Confidence Values”, re-rooting algorithms correctly shift support values to the corresponding branches. Prior to v. 0.9911, there was a minor issue in displaying these values on screen. This was fixed after we contacted the developers. Archaeopteryx does not support the comment notation (e.g., tree *T_C_*).

**ATV** is the predecessor to Archaeopteryx. Different versions seem to alternate between the two possible interpretations of inner node labels. The one we tested uses the branch interpretation of node labels and thus correctly re-roots.

**Dendroscope** versions prior to v. 3.5.0 only offered the node labels as node labels interpretation for our test trees. This lead to incorrect results when re-rooting trees with node labels that actually represented branch support values. Only if the tree also contains branch lengths, Dendroscope interpreted the Newick comments as support values (e.g., tree *T_C_* plus branch lengths). The alternative notation using inner node labels (e.g., tree *T_N_*) is not affected by this and always applies the node label interpretation. This behavior was not fully documented in the manual. We assess the impact of this behavior on published empirical phylogenetic studies in Section “Impact on Empirical Phylogenetic Studies”. In the latest versions of Dendroscope (v. 3.5.0 up to v. 3.5.4), all of our recommendations (see Section “Conclusions”) made in the first bioRxiv preprint [32] of this review were implemented by Daniel Huson. When reading a Newick file with node labels, Dendroscope now explicitly asks the user for the intended interpretation. It also has a menu option to chose between the interpretations.

**ETE (GUI)** [15], [33] is another viewer that supports both interpretations. When reading a Newick formatted tree, it offers an option for specifying label semantics. The comment notation is not supported (e.g., tree *T_C_*).

**EvolView** is able to display numerical values at inner nodes. Re-rooting however misplaces those values to wrong nodes and sets some of them to zero. Re-rooting a given tree several times at different branches results in all inner node values becoming zero. Furthermore, re-rooting does not resolve the initial trifurcation properly, so that the resulting tree contains a multifurcation at node R. The developers are aware of these issues, and intend to fix them in a future release.

**FigTree** is able to display multiple inner node labels using both semantic interpretations. When re-rooting the tree, however, there is no option to define the interpretation of the node labels, that is, FigTree internally always assumes the branch interpretation. Thus, after re-rooting actual node labels, the labels are mapped to wrong nodes. In addition, it can not parse certain Newick variants, such as trees that contain both branch lengths and support values stored as comments.

**iTOL** [18], [34] works correctly. If the inner values are numbers, it implicitly applies the branch support values interpretation. If they are strings, they are interpreted as inner node names. In both cases, re-rooting works as expected. However, it does not offer an explicit option to change this behavior, that is, to interpret numbers as belonging to the nodes, or strings as belonging to the branches.

**PhyloWidget** interprets node labels as node labels. Thus, re-rooting a tree with branch support values yields errors. Also, re-rooting does not resolve the initial trifurcation, similar to EvolView. Phylowidget is no longer maintained, thus its authors recommend not to use it for re-rooting phylogenies or displaying branch support values. Therefore, it is marked as not correct in Table II.

**TreeView** interprets node labels as branch support values and correctly re-roots under this interpretation. However, it displays the values next to the nodes instead of the branches, which may lead to potential confusion.

**T-REX** also applies the branch interpretation and correctly re-roots. The branch support values are however always displayed as percentages, that is, followed by a “%” sign. This is not always the correct or desired way for displaying branch support values. The developers plan to fix this in the next release. Hence, we marked it as almost correct in Table II. T-REX does not work with the comment notation.

#### 2 Bioinformatics Toolkits

**APE** interprets inner node labels as node attributes when re-rooting. We reported this issue to the project maintainers and a new version of the package (v. 3.6) is now available that allows handling node labels as support values when rooting. In addition, a workaround solution is provided in the supplementary material of this manuscript that patches previous APE versions.

**BioPerl** offers options to explicitly load node labels as branch or node attributes. When the branch interpretation is selected, re-rooting algorithms work correctly.

**BioPython,** with the BioPhylo module for handling trees [35], interprets inner node labels as confidence values when parsing a Newick tree. However, those values are handled as node attributes rather than as branch attributes when re-rooting the tree, therefore leading to incorrect positions of the support values. The same behavior is observed when explicitly loading support values using the PhyloXML format. This is currently a known issue in the project and a fix is being developed.

**Dendropy** loads inner node labels as node attributes. Therefore, if those labels are meant to represent support values, re-rooting will lead to incorrect results. The Dendropy documentation explains this behavior in detail, and a workaround is available that permits to re-root trees where bootstrap values are encoded as node labels in the Newick format. A new option has been added in version 4.2 that allows to automatically translate node labels into branch support values when loading a Newick tree, so re-rooting algorithms can be safely applied without further tree processing.

**ETE (API)** [15], [33] offers the same options as when used for tree visualization (see above). Node labels can be loaded as node names (node attributes) or branch support values (branch attributes). When re-rooting, branch support values will be correctly re-mapped to branches.

**Geneious** is able to read both Newick notations, and by default interprets the values as node labels. The branch interpretation is available as an undocumented feature, depending on the naming of those values. However, when re-rooting the tree, the values are treated as belonging to the branches in both cases. This results in misplaced node labels. The maintainers are planning to fix this and to make the interpretation choice more apparent.

**MEGA** [27], [36]–[38] is able to read both notations, and interprets the values as branch support values in both cases. Re-rooting works correctly under this interpretation.

**Mesquite** understands the node label notation, but not the comment notation. By default, it interprets node labels as node labels and correctly re-roots. There is also a function to reinterpret internal node labels and turn them into branch values; re-rooting works correctly after this transformation. For a future release, the maintainers plan to implement a user prompt for choosing the interpretation when a tree with inner node labels is loaded.

**Newick Utilities** does not handle node labels as branch attributes by default, therefore leading to incorrect results when re-rooting Newick trees. After reporting the issue, a previously undocumented option (-`s`) has been documented that permits to automatically interpret inner node labels as branch support attributes.

**Pycogent** interprets inner node labels as support values by default and those are correctly handled by the rooting functions.

### C. Impact on Empirical Phylogenetic Studies

Users, who are not aware of the implicit semantic assumptions of tree manipulation tools, might obtain tree visualizations with incorrectly mapped support values. This is particularly the case if the node interpretation is wrongly applied to branch support values. Most prominently, older versions of Dendroscope (before version 3.5.0, see Section “Results”) implicitly interpret node labels as, simply that, node labels. The extent to which this affects published phylogenies is hard to quantify. This is because all visualized phylogenies in all published papers citing Dendroscope (over 1,200 for the two Dendroscope papers based on Google scholar, accessed on 2016-08-15) would need to be checked and all original tree files would need to be available, which they should be, in principle. Hence, this is also an issue of reproducibility of scientific results—even if in our case it simply boils down to making available a published Newick tree with support values for download. To at least get a feeling of the visualization and reproducibility issue, we contacted the authors of 14 papers that used Dendroscope to visualize trees with support values, published in journals such as Nature, PLOS, BMC, and JBC. Out of the contacted authors, 5 replied, but only two were finally able to provide us with the trees that were used to generate the visualizations in their publications.

In the following, we analyze the trees visualized for these two papers with respect to the correctness of the branch support value mapping.

The first article [39] presents a phylogeny of 80 Arabidopsis accessions (see Fig. 4(b) of [39]) along with bootstrap values for some of the branches. The tree and bootstrap values were inferred with **RAxML** 7.3.5 [40], which writes a tree file that uses Newick comments for storing support values. Dendroscope [12] was used to re-root and visualize the tree. As already mentioned, the tool is able to correctly handle this variant of stored support values. Thus, the error did not occur in this paper and the tree is correctly visualized.

The second article [41] presents several phylogenies for all three domains of life. The trees were inferred using **RAxML** v7.2.6 [40], [42], [43] and **PHYML** v3.0 [7], [44],[45]. Branch support values were estimated with **PHYML** using the SH-like likelihood ratio test [4], which reports support values as node labels. All trees in Figures 2 and 4–7 of [41] were re-rooted using Dendroscope such that they can be more easily compared to the comprehensive trees presented in Figure 1 of the article. In all cases, branch support values were mapped incorrectly to the re-rooted trees in these Figures.

**Figure 2:**
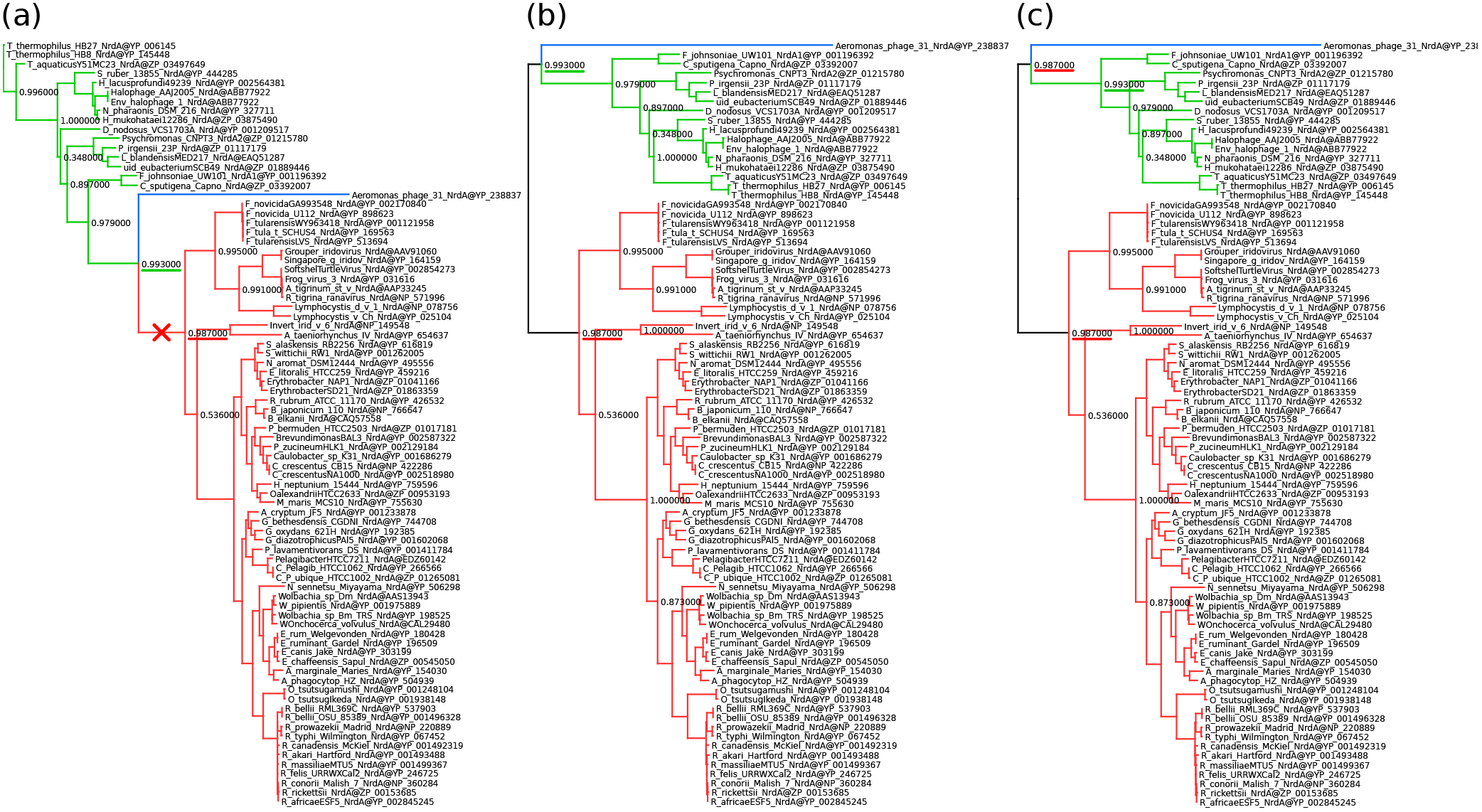
Example of a published phylogeny showing that the issue occurred in real-life data. We used the original data from [41] to recreate Figure 2(a) of [41]. (a) The original tree with the branch used for re-rooting marked by a red cross. (b) The re-rooted tree with incorrectly placed branch support values (e.g., the one underlined in green). This tree was created using Dendroscope 3.4.0. (c) The same re-rooted tree, this time using the updated Dendroscope 3.5.3. The error does not occur, because the correct interpretation of the values was selected. Note that, the value underlined in red is now correctly duplicated at both ends of the root branch. We colored the subtrees to highlight their positions after re-rooting.

We illustrate this in Figure 2. Sub-figure (a) is the original Newick tree used to generate Figure 2(a) in [41]. We have marked the branch used for (re-)rooting the tree by a red cross. We colored the sub-trees so that their corresponding position in the re-rooted tree is easily visible. Sub-figure (b) shows the re-rooted tree using Dendroscope v. 3.4.0, which is identical to the one presented in [41]. The branch support values between the old and the new root node in our Figure 2 are not mapped to the same bipartition in sub-figure

(a) and (b). For example, in sub-figure (a) the support value underlined in green refers to the bipartition `green taxa | blue taxon, red taxa` whereas in sub-figure (b) it refers to the bipartition `red taxa | green taxa, blue taxon`. Fortunately, in this specific case, the incorrectly mapped support values do not change the conclusions of the paper (pers. comm. with Daniel Lundin on 2015-12-28). In sub-figure (c) of Figure 2, we show the correctly re-rooted tree, created with the updated Dendroscope version 3.5.3. The value underlined in green now refers to the correct bipartition. Furthermore, the value underlined in red is correctly duplicated at both outgoing branches of the root.

## III. Conclusions

Our results indicate that an explicit convention and explicit semantics for interpreting node and branch values in tree viewers and other common bioinformatics tools are clearly missing. From the tested viewers, only three (Archaeopteryx, ETE, and Dendroscope from v. 3.5.0 onwards) offer a user dialog to define the semantics of node labels. Older versions of Dendroscope offer an implicit choice depending on the input format. Other viewers can not read certain Newick variants (e.g., Tree *T_C_*). Similarly, bioinformatics toolkits differ in the way node labels are interpreted. Six out of the ten tested toolkits did not provide explicit options for interpreting node labels as branch values. At present, APE, Dendropy and Newick Utilities have now included options for automatically interpreting node labels as branch values when reading and re-rooting trees.

Overall, the tools treat node labels and branch values in their own, often undocumented and implicit, ways. Users must therefore be aware and simply accept the implicit interpretation a particular tool implements.

Furthermore, programs that can infer branch support values use a plethora of distinct output formats. Developers of phylogenetic inference programs may consider storing branch support values using explicit tags as supported by formats such as Extended Newick or PhyloXML [13]. PhyloXML trees are, however, more difficult to parse and yield substantially larger tree files. For instance, our test tree *T_N_* requires 24 bytes in Newick, but 856 bytes in PhyloXML format. Another exemplary 512 taxon tree with branch lengths requires 40,303 bytes in Newick and 239,795 bytes in PhyloXML.

In order to resolve the ambiguity of inner node labels in the Newick format, we recommend to use the comment notation with square brackets to store branch values. This way, the semantics of inner node labels are not overloaded. This is also the variant required by the Nexus standard[46]. Nexus is a container format that internally stores trees in Newick format; in its specification, it refines the original Newick format. However, as this notation “misuses” comments to store meta-data, it is also valid for programs to ignore them. It can thus not be expected to work with current tools, which we showed in this review. Furthermore, particularly when using the comment notation, it is important to explicitly choose the correct interpretation of the stored values.

To address this general problem, we suggest that all tree viewers and toolkits shall offer an explicit option to choose between the two possible interpretations of node labels. Ideally, users should be forced to define the semantics of their node labels before the tree is displayed or re-rooted by the respective tool. This way, accidentally wrong interpretations are avoided and unaware users will become aware of the semantics of inner node labels.

Finally, we suggest that published phylogenies should be re-assessed, if branch support values were stored as node labels in the original Newick files and trees were manipulated using bioinformatics tools (e.g. if Dendroscope prior to v. 3.5.0 was used for re-rooting and tree visualization).

We conclude with some practical suggestions for users of phylogenetic tree viewing tools.

- Pay attention to the options a tool offers for interpreting node labels in Newick files.
- If available, use the option to set the desired interpretation (e.g., Archaeopteryx, ETE, Dendroscope).
- Ensure that re-rooting represents a valid operation for your type of tree and its associated meta-data.
- Double-check your results, maybe try other tools, or conduct a visual inspection, particularly if the original trees were re-rooted or otherwise manipulated.

The behavior of tools can easily be tested with our example trees *T_N_* and *T_C_* that are available for download at https://github.com/stamatak/tree-viz-issues.

## V. Author’S Contributions

LC collected the data, carried out the experiments on tree viewers and on some of the bioinformatics toolkits, and drafted the manuscript. JHC carried out the experiments on most of the bioinformatics toolkits and helped to draft the manuscript. AS conceived the study, participated in its design as well as coordination, and helped to draft the manuscript. All authors read and approved the final manuscript.

## VI. Acknowledgements

This work was financially supported by the Klaus Tschira Foundation and the European Molecular Biology Laboratory (EMBL).

We wish to thank the authors of the papers described in Section “Impact on Empirical Phylogenetic Studies” for sending us their original tree files: Anthony Poole and Daniel Lundin [41], as well as Artem Pankin and Franziska Turck [39].

Furthermore, we want to particularly thank Daniel Huson. He implemented fixes to Dendroscope [12], [47], [48] according to our suggestions shortly after the biorXiv preprint of this review [32] became available. We highly appreciate his feedback on this review and his positive response to our critique and suggestions.

We also want to thank Christian Zmasek (Archaeopteryx, ATV), Zhenxiang Chen and Wei-Hua Chen (EvolView), Ivica Letunić (iTOL), Gregory Jordan (PhyloWidget), Vladimir Makarenkov (T-REX), Emmanuel Paradis (APE), Chris Fields and Jason Stajich (BioPerl), Peter Cock (BioPython), Jeet Sukumaran and Mark T. Holder (Dendropy), Alexei Drummond and Richard Moir (Geneious), Sudhir Kumar and Koichiro Tamura (Mega), David Maddison and Wayne Maddison (Mesquite), Thomas Junier (Newick Utilities) and Daniel McDonald (PyCogent) for their feedback and discussions regarding this review and their software.

